# Global motion processing by populations of direction-selective retinal ganglion cells

**DOI:** 10.1101/572438

**Authors:** Jon Cafaro, Joel Zylberberg, Greg Field

## Abstract

Simple stimuli have been critical to understanding neural population codes in sensory systems. Yet it remains necessary to determine the extent to which this understanding generalizes to more complex conditions. To explore this problem, we measured how populations of direction-selective ganglion cells (DSGCs) from mouse retina respond to a global motion stimulus with its direction and speed changing dynamically. We then examined the encoding and decoding of motion direction in both individual and populations of DSGCs. Individual cells integrated global motion over ~200 ms, and responses were tuned to direction. However, responses were sparse and broadly tuned, which severely limited decoding performance from small DSGC populations. In contrast, larger populations compensated for response sparsity, enabling decoding with high temporal precision (<100 ms). At these timescales, correlated spiking was minimal and had little impact on decoding performance, unlike results obtained using simpler local motion stimuli decoded over longer timescales. We use these data to define different DSGC population decoding regimes that utilize or mitigate correlated spiking to achieve high spatial versus high temporal resolution.

## Introduction

Sensory systems encode and decode information across populations of neurons. Understanding such population codes is fundamental to understanding the function of neural circuits and sensory processing (Pouget et al., 2000; Panzeri et al., 2015). Population codes are likely optimized for natural sensory stimuli but they are often probed using simple and artificial stimuli (Felsen et al., 2005; Fitzgerald and Clark, 2015). Such simplifications may limit an understanding of population codes and neural function in ethological contexts. In this paper, we examine a canonical population code, direction coding in mammalian ON-OFF (oo)DSGCs, in the context of global motion of a natural scene.

In the mammalian retina, there are four types of ooDSGCs, each tiling space with their dendritic and receptive fields (Barlow et al., 1964; Devries and Baylor, 1997; Demb, 2007; Vaney et al., 2012; Morrie and Feller, 2016). These types differ primarily in their preferred direction of motion, which are organized along four cardinal axes (Oyster and Barlow, 1967; Vaney, 1994; Kay et al., 2011; Trenholm et al., 2013; Yao et al., 2018). Direction is encoded across the four types by their relative firing rates. This produces a population code for direction that is relatively invariant to object speed and contrast (Nowak et al., 2011; Zylberberg et al., 2016). ooDSGCs have been largely considered responsible for signaling local motion, because global motion attenuates (but does not eliminate) their responses (Vaney et al., 2001; Chiao and Masland, 2003; Olveczky et al., 2003; Hoggarth et al., 2015). A separate class of DSGCs, so-called ON DSGCs (oDSGCs), are minimally attenuated by global motion, and have thus been assumed to play a dominant role in signaling global motion (Oyster, 1968; Simpson et al., 1988). Correspondingly, previous studies examining the fidelity and accuracy of the ooDSGC population code have focused on local motion and artificial stimuli that are decoded at relatively long timescales (Fiscella et al., 2015; Zylberberg et al., 2016). These studies largely pointed toward a high-fidelity code that utilizes correlated activity in nearby ooDSGCs to signal the direction of local motion. However, recent work indicates that ooDSGCs may be organized to encode self-motion, a global motion signal (Kay et al., 2011; Dhande et al., 2013; Sabbah et al., 2017). This motivates an examination of ooDSGC individual and population responses under conditions in which the stimulus is a natural scene moving globally and dynamically on the retina. It also motivates understanding how the direction of global motion can be decoded from populations of mammalian DSGCs and the extent to which concepts applicable to decoding local motion at long timescales apply to decoding global motion at shorter, and perhaps more behaviorally-relevant, timescales.

To study DSGC responses, we recorded simultaneously the spiking activity from hundreds of retinal ganglion cells (RGCs) using a large-scale multielectrode array (MEA). We distinguished DSGCs from other RGCs based on their responses to drifting gratings (Elstrott et al., 2008; Yao et al., 2018). We then projected dynamically moving natural images onto the retina: the motion is ‘dynamic’ because the direction and speed are not constant. Individual ooDSGCs and oDSGCs exhibited similar encoding of dynamic global motion stimuli: they both integrated and low-pass filtered direction signals over a timescale of ~200 ms; they were both broadly tuned; and they both exhibited similar spike rates. Importantly, both ooDSGC and oDSGCs exhibited little trial-to-trial variability in their responses to dynamic global motion, indicating that while the responses were sparse, they were reliable.

We then utilized our more complete populations of ooDSGCs to examine the limitations inherent in decoding dynamic global motion signals from small and large ooDSGC populations. For a local quartet of ooDSGCs (each with a different preferred direction), determining the direction of global motion was marginally better than chance at short timescales (~100 ms). Decoding accuracy was improved by longer temporal integration of ooDSGC signals, however this is only an effective decoding strategy when changes in motion direction are infrequent and when the animal does not need to rapidly respond to a change in the motion signal. When motion direction changes frequently, large populations of ooDSGCs are needed to accurately and rapidly (< 100 ms) decode the direction of global motion. Large populations of ooDSGCs are available for decoding at no cost to spatial resolution because the nature of the motion signal is global. Furthermore, the short integration times used when decoding large populations result in largely uncorrelated population activity, which is counter to previous results decoding local motion at long timescales (Franke et al., 2016; Zylberberg et al., 2016). This limits the impact of correlated spiking on decoding accuracy in a dynamic global motion context. Thus, large populations of nearly-independent ooDSGC signals integrated over short timescales enables rapid decoding of direction. We generalize these findings to illustrate the tradeoffs inherent in decoding visual signals that vary in space versus time.

## Results

Visually driven responses of retinal ganglion cells (RGCs) were measured ex vivo using a multi-electrode array (MEA) (Elstrott et al., 2008; Yao et al., 2018). Responses to drifting gratings distinguished ooDSGCs and oDSGCs from other RGCs over the MEA (see Methods). To measure the responses of DSGCs to dynamic global motion (Fig 1A), a natural scene from the Van Hateren image database (van Hateren and van der Schaaf, 1998) was dynamically moved over the retina. This paradigm drove the responses of dozens of identified and simultaneously recorded ooDSGCs and oDSGCs.

**Figure 1:**
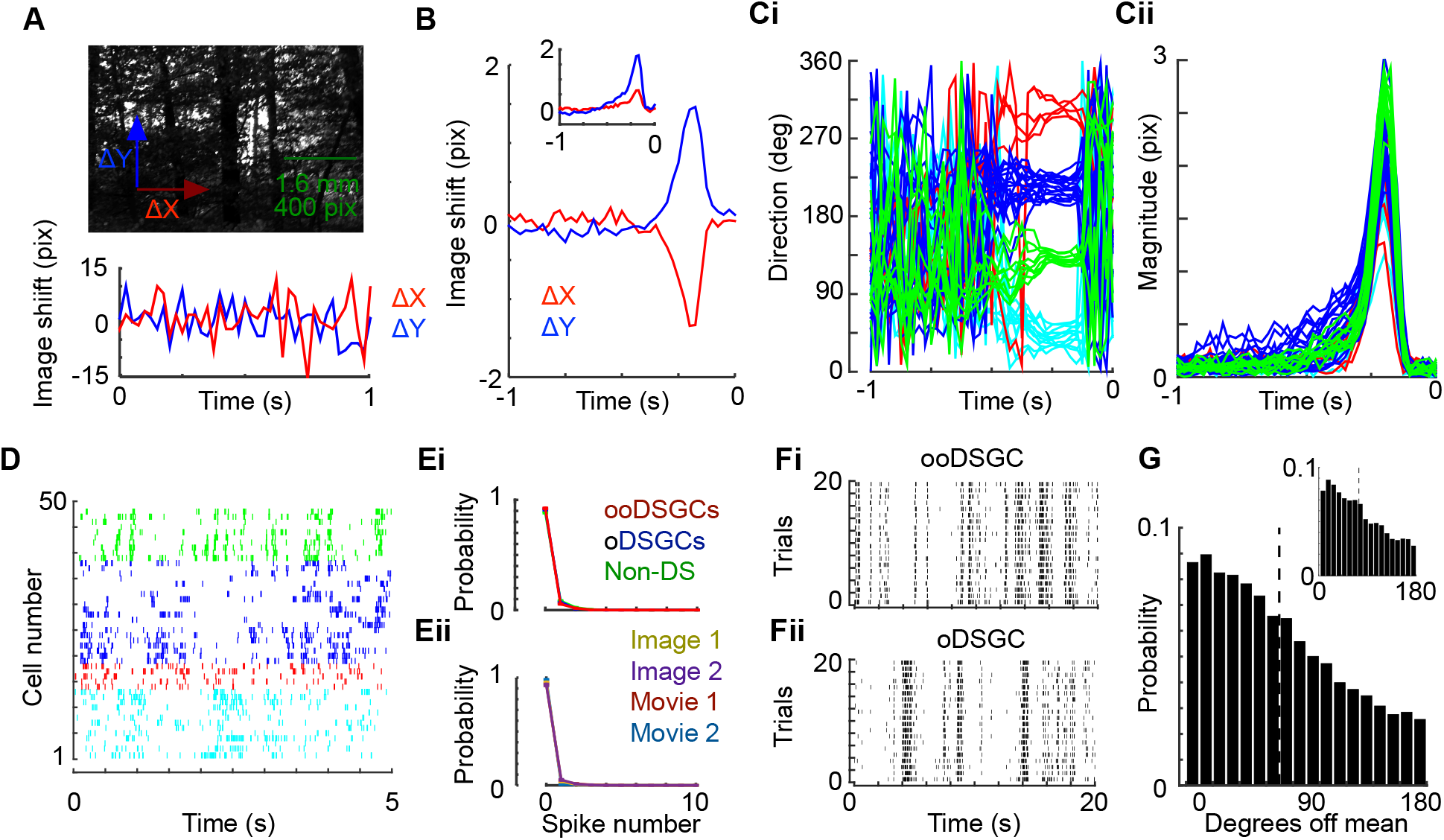
DSGCs integrate the direction of global motion over time and respond sparsely with broad tuning to natural images. **A.** Natural image presented (top) and displaced according to ΔX and ΔY (bottom). **B.** Spike triggered average of ΔX and ΔY for single example ooDSGC and an oDSGC (inset). **C.** Direction (Ci) and magnitude (Cii) of motion calculated from spike triggered average (STA) for all ooDSGCs in a single retinal recording. Color coded according to preferred direction in grating stimulus. **D.** Spike times during five seconds of dynamic global motion stimulus in (e.g. A) for all ooDSGCs in a single retinal recording; color indicates preferred direction determined form a drifting grating. **E.** Probability distributions for DSGCs in a single retinal recording using a jittered image (Ei) and for ooDSGC and oDSGCs in 2 different retinal recordings using 2 different jittered images and 2 different natural movies (Eii). Movies included video from a camera mounted on a mouse (Movie 1) and a cat (Movie 2). **F.** Spike raster of a single ooDSGC (Fi) or oDSGC (Fii) simultaneously recorded over several repeated presentations of the same jittered image. **G.** Difference between preferred direction from STA (panel Ci) at time of peak magnitude (panel Cii) and direction preceding each spike. Histogram includes data from all ooDSGC cells and all spikes from a single retina (n = 49); similar results were observed in a second retinal recording. The inset shows the same analysis for all oDSGCs in the same recording.

### Individual ooDSGCs encode direction via integration of dynamic global motion

Examining responses during direction changes can reveal how DSGCs integrate over motion direction history - an important factor in their response to global motion of a natural scene. Thus, we begin by focusing on ooDSGCs, and analyzing the relationship between their spiking and dynamic global motion of a natural scene. We randomly and iteratively translated a natural scene on the retina while recording ooDSGC spikes (see Methods). The, X and Y positions of the image were shifted in each frame of the video display by ΔX(t) and ΔY(t). Image position shifts were sampled independently from a Gaussian distribution, generating an approximately ‘white noise’ motion stimulus (Fig 1A; see Methods). The image shift distribution had a zero mean with a standard deviation ~20 μm/frame (~25 deg/s). This value was chosen to maximize responses from ooDSGCs and fall within the range of eye movement velocities in freely moving rats (Wallace et al., 2013) and retinal image motion in rabbits (Van der Steen and Collewijn, 1984). We calculated the correlation between ΔX and ΔY values and spike rate, yielding a spike-triggered average (STA) of the displacements in X and Y for each cell (Fig 1B)(Borghuis et al., 2003; Perge et al., 2005; Kuhn and Gollisch, 2019). Translating these Cartesian to polar coordinates facilitated visualizing the STA-directions of all ooDSGCs simultaneously (Fig 1C). Approximately 500 ms preceding a spike, the average motion direction fluctuated randomly for every ooDSGC (Fig 1Ci). However, between 300 to 100 ms preceding a spike, the motion direction coalesced to one of four cardinal directions. These results indicate that each ooDSGC encodes global motion along one of four directions and that spiking depends on the motion direction over ~200 ms temporal window, with ~100 ms latency to spiking (Fig 1C).

There are two possible strategies by which ooDSGCs may encode this motion. First, ooDSGCs may simply integrate motion signals over a temporal window. Alternatively, they may signal a change in direction by differentiating the motion trajectory. Differentiation is a common computation performed by the receptive fields of most RGCs (e.g. center-surround antagonism (Kuffler, 1953; Perge et al., 2005; Schwartz et al., 2007)). Pure integration requires a monophasic dependence on motion trajectories preceding spikes, while differentiation (in the case of direction changes >90 degrees) requires a biphasic dependence on motion trajectories. Every ooDSGC exhibited a monophasic direction STA (peak/trough ratios at 14.2 ± 2.7; Fig 1B), with a mean half width of 111 ± 2 ms (Fig 1Cii). Thus, ooDSGCs encoding appears more related to the integration of direction for global motion stimuli within relatively short time windows preceding their spikes; they do not appear to explicitly signal changes (differentiation) in the motion direction. Below we explore the implications of needing to decode ooDSGC signals that are updated continuously over short (200 ms) time windows.

### Individual ooDSGCs generate sparse and broadly tuned responses to naturalistic global motion

The analyses above reveal the average motion kinetics and directions that precede ooDSGC spiking for global motion in a natural scene. However, the fidelity of encoding, and the accuracy of decoding, will depend strongly on the spiking dynamics elicited by these stimuli. Spiking was infrequent in ooDSGCs to natural scene global motion (Fig 1D,E), consistent with other measures of RGC activity during natural movie presentations (Koch et al., 2006). For the global motion stimulus, firing rates ranged from 0.8 to 8.5 Hz, with one or more spikes occurring in a single neuron on less than 8% of the video frames (40 Hz frame rate). This result was replicated with several different images and natural movies from cameras that were head mounted to animals (Fig 1Eii; See Methods), demonstrating that global motion in natural scenes typically evokes sparse responses across ooDSGCs.

One question that arises is whether or not these stimuli were reliably driving spikes in ooDSGCs, given the low spike rates. Repeated presentations of the same stimulus produced stereotyped ooDSGC responses (Fig 1Fi), indicating that the response sparsity is not simply a result of presenting a stimulus that is incapable of evoking a response. Instead, these stimuli generated sparse responses that were reliable from trial to trial – within each response frame the spike count mean was approximately equal to the variance (mean fano factor ± SEM = 1.1 ± 0.05). However, the motion direction ~200 ms preceding individual spikes was highly variable (Fig 1G). To quantify the variability in motion direction preceding spikes, we calculated the difference between the direction of motion preceding each spike and the STA direction (evaluated at the peak of the STA magnitude). This distribution is broad and on average the direction preceding a spike differs from the mean (preferred) direction by ~70 degrees (Fig 1G, dashed line). This variability will limit decoding performance, because the presence of a spike poorly constrains the preceding motion direction.

Variability in the pre-spike direction likely reflects several sources including: the tuning width of the ooDSGC, different direction trajectories across video frames filling the ooDSGC integration time, and aperture effects that allow local orientation to influence apparent direction within a receptive field (McDermott et al., 2001; Sung et al., 2009; Kane et al., 2011). Irrespective of the source, the stimulus variability preceding ooDSGC spiking combined with infrequent spiking, will limit the accuracy with which direction of global motion can be decoded from ooDSGC populations. Below we assess if the response properties described above are unique to ooDSGCs, or whether these observations also apply to oDSGCs.

### oDSGCs respond similarly to ooDSGCs

Previous work has suggested that signaling self-motion is performed by oDSGCs while ooDSGCs signal local object motion (Vaney et al., 2001). Thus, we compared the responses of oDSGCs to ooDSGCs to see if they exhibited distinct response properties to global motion in natural scenes. First, oDSGCs showed similar monophasic temporal integration to ooDSGCs (Fig 1B inset). Second, oDSGCs showed similar response sparsity to the same global motion stimuli (Fig 1Ei). Indeed, all recorded RGCs exhibited similar response sparsity (Fig 1Ei). oDSGCs also exhibit similarly reliable responses to repeated presentations of the same global motion sequence for a natural scene (Fig Fii) and similar direction variability preceding a spike as ooDSGCs (Fig 1G, inset). Thus, we did not observe clear differences in the response statistics or encoding properties between oDSGCs and ooDSGCs to global motion of a natural image.

The analyses below leverage the simultaneously recorded populations of ooDSGCs to test the ability to decode the direction of global motion from those populations and analyzes the factors limiting the accuracy of that decoding.

### Quartets of ooDSGCs exhibit limited accuracy in signaling the direction of global motion

To begin to understand how the response properties of ooDSGC impact the decoding of motion, we applied an optimal linear estimator (OLE) to the responses from quartets of simultaneously recorded ooDSGCs. In brief, an OLE assigns a set of weights to each cell which, when scaled by the response of that cell and summed across cells, will minimize the mean squared error of the prediction (Fig 2A; see Methods). Each quartet consisted of ooDSGCs with different preferred directions and cells within 200 μm of one another (Fig 2B). We begin with quartets of ooDSGCs because they form an elementary unit of a population code. Specifically, spikes from one ooDSGC poorly constrain motion direction, because of the broad direction range that can precede a spike. However, spikes distributed across a quartet of ooDSGCs can, in principle, be used to more accurately decode motion direction (Georgopoulos et al., 1986). We begin with an OLE because it is a simple decoder that performs nearly optimally on ooDSGC population responses and can be simply implemented by downstream neurons (Salinas and Abbott, 1994; Fiscella et al., 2015).

**Figure 2:**
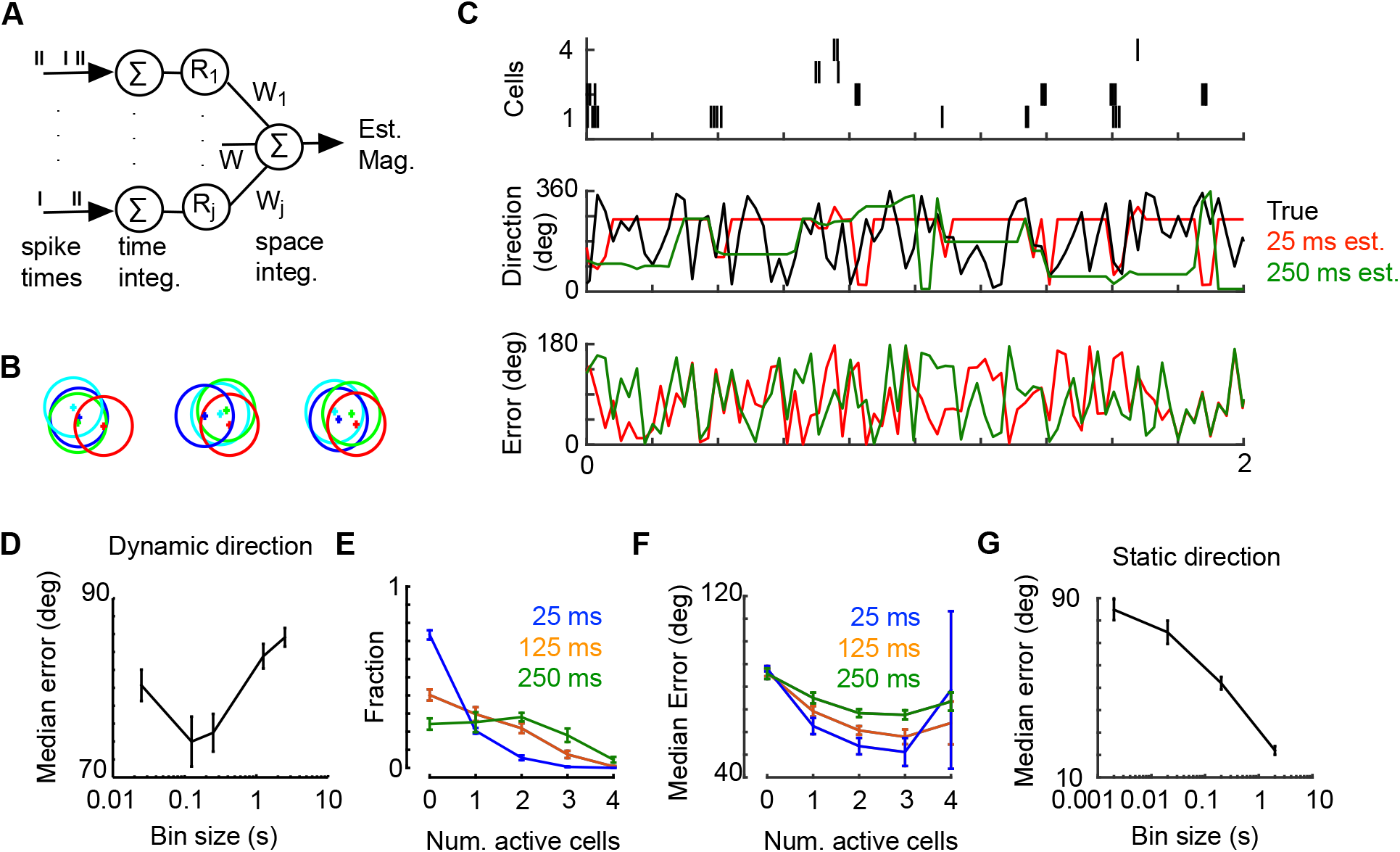
ooDSGC quartets are limited in their decoding accuracy by response sparsity and dynamic motion. **A.** Schematic of the decoder, the optimal linear estimator (OLE). **B.** Examples of four ooDSGC quartets. Relative location of each cell is marked from position on array and circles drawn, 300 μm diameter, provide a scale bar near the size of ooDSGC dendritic fields. **C.** *Top:* Spike response from an example quartet during a dynamic global motion stimulus (spike times are uniformly shifted for optimal decoding). *Middle:* Decoder provides estimate of direction for each frame change using two different spike integration times. *Bottom:* Acute difference between OLE and actual direction for each integration time. **D.** Median error over all decoded time points as a function of integration time from one retina (N=15 quartets). **E.** Fraction of bins with non-zero spikes in 0-4 cells in each quartet (same retina as in D). **F.** Median error as a function of number of cells in quartet with non-zero spikes (same retina as in D). **G.** Median error for all decoded time points as a function of integration time when decoding drifting image with direction and speed held constant (see Methods). Separately recorded retina from panels B-F (7 quartets). Error bars (D-G) show standard deviation across quartets.

The first question we address with this approach is, ‘how accurately can global motion in a natural scene be decoded from the responses produced by a local quartet of ooDSGCs?’ The answer is likely to depend on the duration over which the decoder integrates signals from the ooDSGCs, and the dynamics of the motion (e.g. how frequently the velocity changes). First, we examined the dependence on integration time. When integration time is short, the decoder is forced to estimate direction from responses produced within single video frames (~25 ms). This yielded low accuracy estimates of motion direction; the median expected error was ~80 degrees (Fig 2C-D). Note this analysis allows for a latency between the stimulus direction and ooDSGC responses (see Methods). The median error is reported throughout and provides the minimum error in decoding 50% of the time bins, an appropriate quantity when decoding continuously. For comparison, chance performance in direction estimation would be 90 degrees, and perfect performance would be 0 degrees. A major contributor to this high uncertainty in motion direction is that within ~25 ms, the most frequent output from the quartet of ooDSGCs is zero spikes (Fig 2E). When there are no spikes, the decoder assigns a default constant, effectively guessing at the direction of motion. It is worth noting, that in a stimulus regime with constantly changing direction, this default is no worse than assuming the direction that was last decoded when spikes occurred.

To test that the high error at short integration times results from the sparsity of the population response, we analyzed the frequency with which a given number of ooDSGCs responded within a quartet. For short integration times there is a high probability of zero spikes from any ooDSGC in the quartet (Fig 2E). Furthermore, decoding error depended on the number of cells responding within a given integration window - the error decreased sub-linearly for increasing cell numbers (Fig 2F). Errors were high when four cells were responding in the same bin, which results from cancelation of oppositely tuned neurons.

One path toward improving decoding performance is for the decoder to integrate over longer time windows. This would allow for a larger fraction of decoded epochs to contain at least one spike from the quartet of ooDSGCs. However, increasing the integration time to 125 ms (5 stimulus frames) only modestly decreased the error of direction estimates to ~74 degrees. Furthermore, for longer integration times, average direction error increased (Fig 2D). Thus, decoding global motion from local quartets of ooDSGCs exhibits limited accuracy.

The increase in decoding error at longer integration times is likely a result of the dynamic stimulus, which frequently changes directions. Thus, integrating for longer periods of time incurs a cost: the inability to decode frequent changes in direction. To test this hypothesis, we switched from decoding an image that changed direction and speed dynamically to a drifting natural image that moved in a constant direction and speed (see Methods). As hypothesized, images moved in a static direction show only increases in accuracy with increasing integration time (Fig 2G), as the decoder was afforded the opportunity of accumulating spikes over long periods of time without a change in direction. Using a 2 s integration window to decode the direction of a drifting natural image reduced the average error down to ~20 degrees when decoding from a quartet of ooDSGCs.

The analyses above show that quartets of local ooDSGC provide little information about global motion direction in a natural scene at short time scales. Their limited decoding accuracy is largely due to the sparse (infrequent) spiking generated by the stimulus. Furthermore, decoding is limited to short integration times when motion is dynamic because integrating over longer time windows fails to track changes in motion direction. This is at least partly a consequence of ooDSGCs integrating, instead of differentiating, motion (Fig 1B). If decoding accuracy is limited by the sparsity of the population response, do larger populations of ooDSGCs allow for more accurate decoding of dynamic motion at short integration times?

### Large ooDSGC populations can encode direction continuously over short time scales

To begin to test the effect of ooDSGC population size on decoding global motion, we decoded the direction of dynamic global motion using the responses of all ooDSGCs measured in an experiment (Fig 3A). While these populations are not complete, due to imperfect sampling of RGCs over the MEA, this analysis permitted data-based decoding on 43-50 ooDSGCs in individual experiments. Furthermore, the population spanned lengths of ~750 um (25° of visual arc) on the retina.

**Figure 3:**
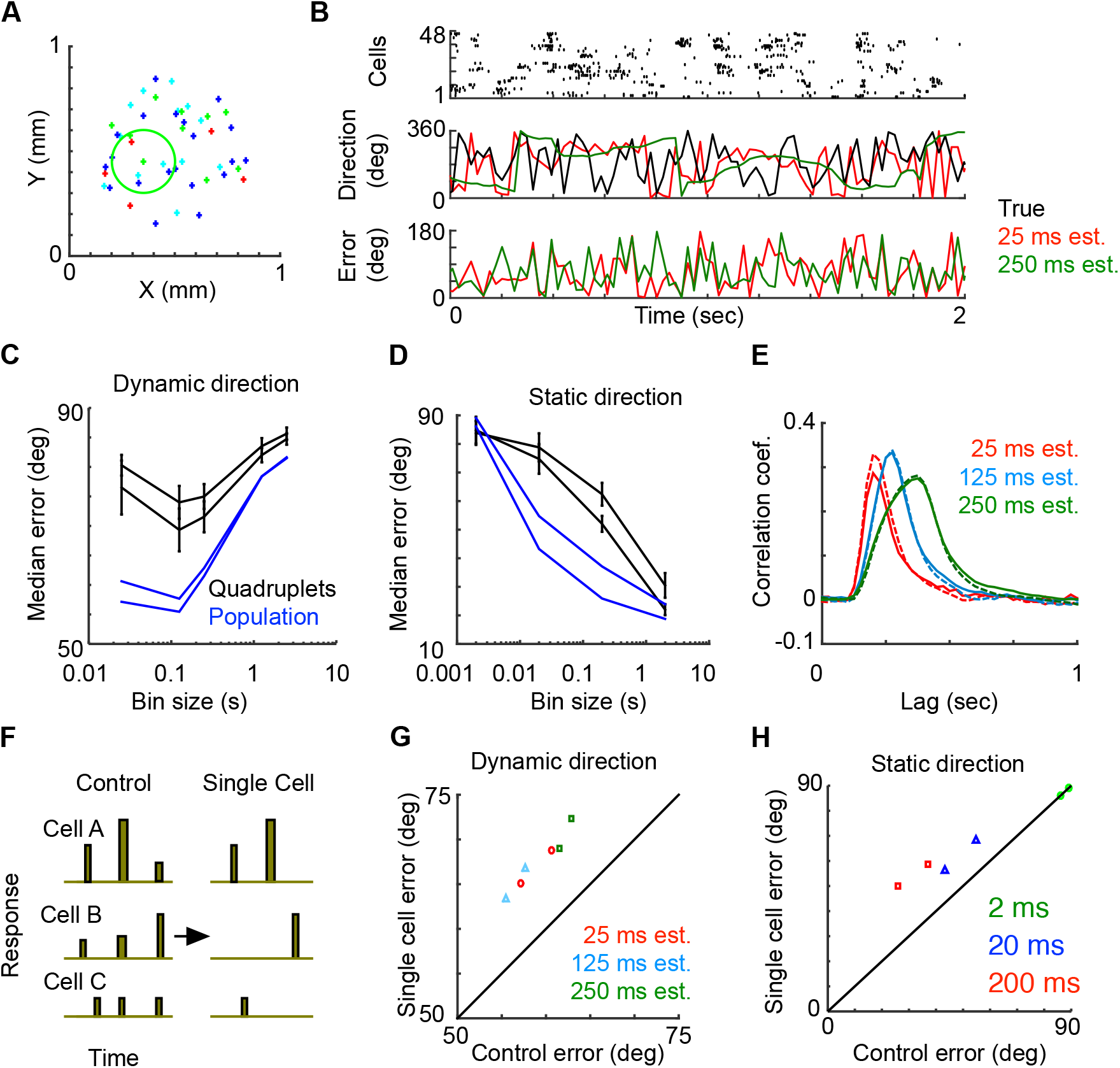
Larger populations of ooDSGCs improve decoding accuracy with shorter integration times. **A.** Location on array of all ooDSGCs recorded in single retina. Circle provides estimate of the size of receptive field and colors indicate preferred direction. **B.** *Top:* Spike response from recorded ooDSGC population during dynamic global motion stimulus. *Middle:* Decoder provides estimate of direction for each frame change using two different spike integration times. *Bottom:* Acute difference between OLE and actual direction for each integration time. **C.** Median error for all decoded time points as a function of integration time. For both the quartets (as in Figure 2D) recorded in 2 different retinas and the entire population for 2 different retinas. No error bars are provided for the estimate across the entire population. **D.** As in panel C using an image with a constant direction (see Methods). **E.** Cross correlation between stimulus image displacement estimate and actual stimulus (ΔX=solid or ΔY=dashed line). **F.** Schematic showing “single cell” manipulation of input to OLE. **G.** Median error using control and “single cell” input to OLE. Results from two retinas are shown at 3 different spike integration times. **H.** Same as panel F using image with direction held constant.

Larger populations of ooDSGCs increase the frequency with which one or more cells spike for a given integration time, relative to quartets. This effectively decreases the sparsity of the population response to which the decoder has access. As a result, the median error from decoding these larger population responses was significantly smaller than decoding quartets, particularly for short integration times (Fig 3B-C). For example, at ~25 ms (a single video frame), decoding error was reduced to 55-60 degrees for a population of 48 ooDSGCs, down from 80 degrees for a quartet. It is notable that decoding direction from a population on a single frame was so accurate, given that a single frame is much briefer than the integration time of the STA (Fig 1B).

Similar to the results from ooDSGC quartets, increasing the integration time also caused an increase in median errors for larger ooDSGC populations (Fig 3B-C). This increase is because the global motion is dynamic, causing the decoder to estimate a single direction of motion from responses that are produced by multiple directions. When the direction of the stimulus was constant, longer integration times resulted in a monotonic decrease in error for large populations of ooDSGcs (Fig 3D). For long integration times (2 s), decoding error fell to ~20 degrees with a population of 48 ooDSGCs. Thus, larger ooDSGC populations allow for more accurate decoding of global motion in natural scenes within briefer integration times. However, long integration times limit decoding performance when global motion changes dynamically.

Thus far we have shown that long stimulus integration impairs the ability of ooDSGC populations to accurately estimate dynamic motion. Long stimulus integration has an additional cost, which is to delay the time at which direction estimates are most accurate relative to the stimulus. To measure this delay, we computed the cross correlation between the actual and estimated image displacements (in ΔX and ΔY). The cross-correlation between these values was significantly delayed and broader at longer integration times (Fig 3E). Thus, integration over short timescales allows downstream circuits to decode more rapidly, thereby following more frequent changes in direction. This is only achievable with large populations of ooDSGCs because quartets perform marginally better than chance within the same integration times.

Increasing the population size could improve decoding in two different ways: 1) by increasing the number of time points with single responsive cells; and/or 2) increasing the number of time bins with multiple responsive cells. To measure the extent to which the error depended on a simultaneous multi-cell response, the OLE was trained on the full response set and tested on either the full response set, or on a modified response set in which only a single cell response (the largest response) at each time bin was provided to the decoder (Fig 3F). If direction decoding is entirely mediated by single cells, then there should be no difference between using the full and modified response sets. There was a significant increase in the error when decoding on the modified response set in both the dynamic (Fig 3G) and static (Fig 3H) direction stimuli. Thus, the decoding accuracy in larger populations relies on simultaneous activity from multiple ooDSGCs.

The simultaneous activity between ooDSGCs that underlies a population response could arise purely through independent responses across ooDSGCs or through correlated subsets of ooDSGCs. In the next section we examine the extent to which the accuracy of rapid decoding in large ooDSGCs populations relies on response correlations within the population.

### Rapid-global direction of motion is encoded by large populations of independent ooDSGCs

Natural scenes have local intensity correlations that result in correlated activity between nearby RGCs (Simoncelli and Olshausen, 2001; Pitkow and Meister, 2012). Recent work has indicated that such response correlations promote robust decoding by maintaining the relative activity between ooDSGCs with different preferred directions (Franke et al., 2016; Zylberberg et al., 2016). To what extent are response correlations important to maintaining the accuracy of rapid decoding of global motion from large ooDSGC populations?

To understand how the correlation structure contributes to decoding accuracy, we measured and manipulated response correlations across the ooDSGC populations. In this section we focused entirely on the static direction stimulus, which permitted manipulations that would be impossible across a dynamic direction stimulus.

First, we examined the correlation structure in the population by mapping the pairwise correlation coefficients as a function of (1) distance between pairs, (2) relative preferred direction, and (3) integration time (Fig 4Ai-iii). The correlation coefficients were calculated within a trial and averaged across all trials and directions. Thus, the response correlations reported here include signal and noise correlations and measure the tendency of cells to respond to the same image structure. The correlation coefficient between pairs of ooDSGCs increases with the integration time used to calculate the responses, as previously noted (Cohen and Kohn, 2011), short integration times diminish correlations (Fig 4A-note axes scale). Thus, over short integration times, correlations are small, suggesting they may not influence decoding accuracy to the extent observed in previous studies that considered longer integration times (Franke et al., 2016; Zylberberg et al., 2016).

**Figure 4.**
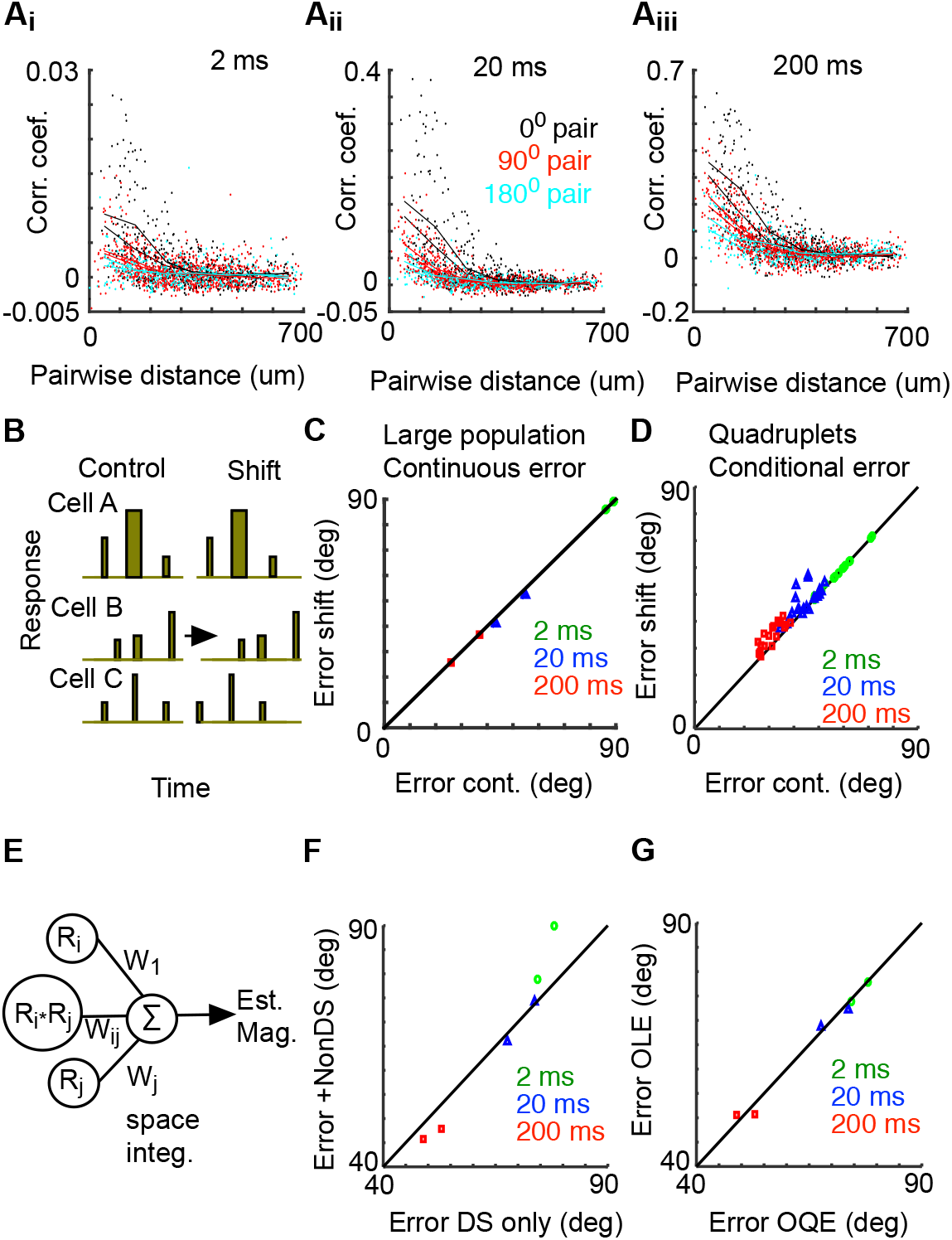
Correlation coefficients are small on short time scales and have little impact on decoding accuracy. **A.** Correlation coefficient between pairs of ooDSGCs as a function of the distance between pairs. All possible pairs within a retina are plotted for two different retinas (points) and the averages within a binned I distance are plotted for each retina (lines). Pairs are color coded by the angle difference of their preferred direction during the drifting grating. Panels i-iii present the same data but calculated using three different integration times. **B.** Schematic illustrating the “shift” manipulation in the input to the OLE. **C.** Median error using control and “shifted” input to the OLE. **D.** Same as C but using ooDSGC quartets and decoding only during the periods of high magnitude direction estimates. **E.** Schematic illustrating decoding with OQE. **F.** Decoding error of small local population of ooDSGCs and Non-DS cell responses using an OQE. **G.** Decoding error of small local population of ooDSGCs only using an OQE and OLE.

However, for a given integration time, the correlation is higher for cells that are spatially closer and modulated less prominently by their relative preferred directions (eg. Fig 4Ai) (Pitkow and Meister, 2012). This reflects the increased tendency of nearby cells to respond to the same part of the image as the dominant determinant of correlation structure. This led us to ask if the higher correlations in nearby cells are important in maintaining the accuracy of decoding from large ooDSGC populations over short time scales. In other words, are responses to global motion encoded by many small-local populations of correlated cells?

To test how decoding error depends on the correlations between ooDSGCs, the OLE was tested on either the unmodified (control) response set or a decorrelated (shifted) response set, in which the response bins were shifted in time during a drifting image (Fig 4B). Shifting responses in time independently across ooDSGCs eliminates correlations due to local contrast fluctuations in the stimulus and noise correlations introduced by retinal circuits. However, this manipulation maintains correlations due to the direction of motion. Thus, shifting responses in time undermines the population response structure caused by the particular spatial locations of the cells, and is similar to selecting populations of ooDSGCs randomly in space. Across a range of integration times, the ‘shifted’ response sets showed little change in continuous decoding error (Fig 4C). This result suggests that decoding of direction from large ooDSGC populations does not depend on correlations, even when those correlations are relatively large.

These result differ substantially from previous findings where trial-to-trial noise correlations alone were shown to significantly decrease decoding error by maintaining orthogonality between signal and noise (Franke et al., 2016; Zylberberg et al., 2016). However, measured populations of incomplete mosaics under sample overlapping groups of cells, which may undermine the impact of large correlations measured during long integration times. To better understand how integration time impacts the decoder’s sensitivity to correlations, we focused on decoding in quadruplets, which are selected based on their overlap, during times periods when they were strongly responding. We used the OLE magnitude to select the time bins in which the ooDSGCs population was responding most strongly (Fig 2A). The OLE magnitude will be highest when multiple cells, with similar tuning are responding most strongly. We assess the median error during the top 10% magnitude responses and term this the “conditional error” (because it is conditioned on the OLE magnitude being high). The conditional error in quadruplets was sensitive to correlation structure at long integration times, increasing the error when correlations were disrupted (Fig 4D), but not at short integration times. Thus, the impact of correlations on decoding depends critically on the integration time – correlations being important when decoding large responses integrated over long time windows. We later extend these results to large modeled DS populations with complete mosaics (Fig 5).

**Figure 5.**
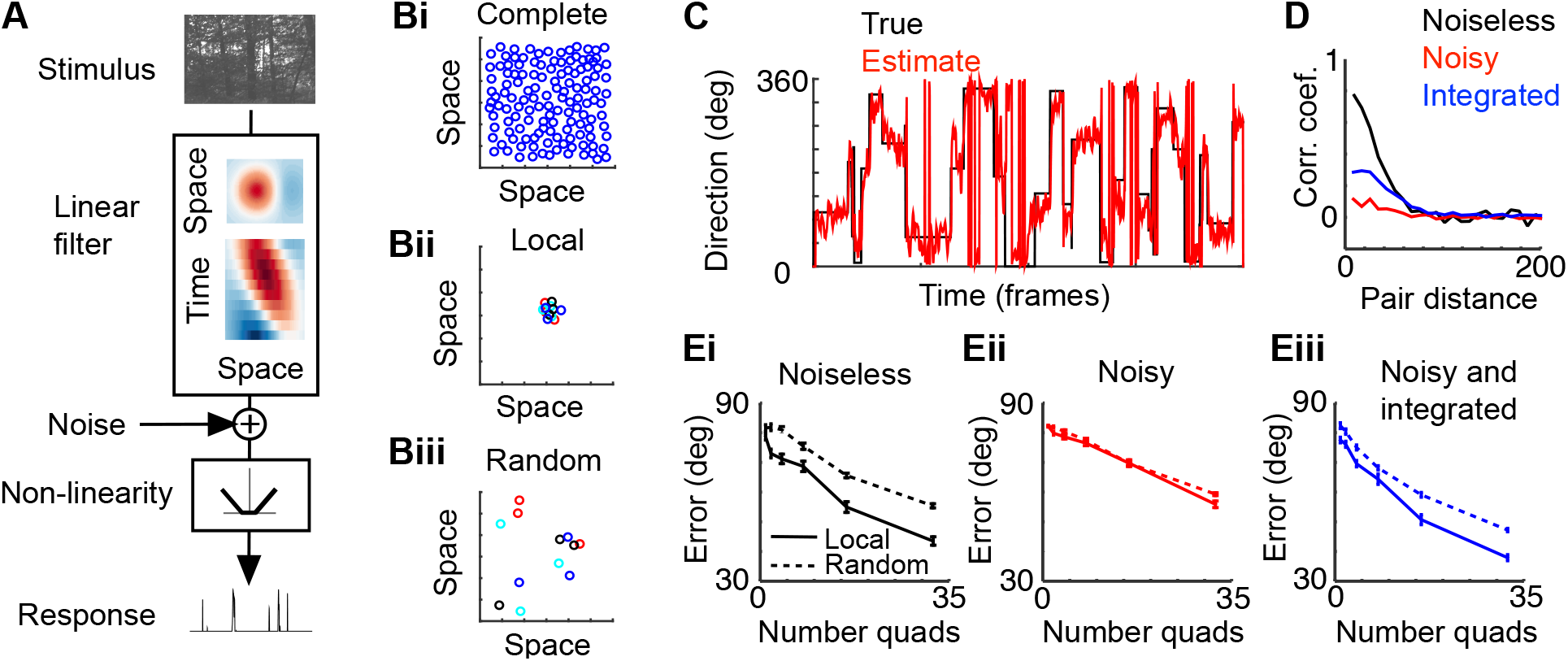
Model of ooDSGC populations demonstrates how natural images, integration time and noise influence correlations and decoding. **A.** Schematic of a single modeled DS-unit. **B.** Example of receptive field positions subsampled from complete DS-unit mosaics (Bi-showing 1 of 4 mosaics) when decoding three spatially local (Bii) or random (Biii) quadruplets. Circles illustrate one standard deviation of receptive field size. **C.** Example of image direction and decoded direction estimate at each point in time. **D.** Average pairwise correlation coefficient within a large population of modeled DS-units as a function of distance between pairs. **E.** Decoding error versus size of DS-unit population in either a noiseless model (Ei), a noisy model (Eii), or a noisy model with temporal integration (Eiii).

### Non-DS cells do not improve direction decoding over short integration times

The analyses above indicate that correlations between ooDSGCs are not useful to account for local image intensity when decoding the direction of motion in short integration times. However, local image intensity influences the spike rates of non-DS, as well as DSGCs. Potentially, non-DS RGCs that share substantial receptive field overlap with ooDSGCs could be used to help decode the direction of motion by discounting local image intensity. Indeed, correlations between tuned and untuned neurons have been shown to improve decoding from other neural populations (Leavitt et al., 2017; Zylberberg, 2018).

To assay if non-DS RGCs can help decode the direction of motion we decoded the direction of motion using both identified ooDSGCs and non-DS cells. Using local groups with substantial receptive field overlap (~30 neurons), the direction of motion was first decoded using an OLE. There was not a significant difference between decoding performance with or without non-DS cells (data not shown). This result is expected, because an OLE, while sensitive to correlation structure, does not explicitly use correlated activity to decode (Schneidman et al., 2003; Shamir and Sompolinsky, 2004). Alternatively, an Optimal Quadratic Estimator (OQE) explicitly uses correlations between neurons to decode by weighting the synchronous activity between neurons to estimate the direction of motion (Shamir and Sompolinsky, 2004) (Fig 4E; see Methods). The OQE accuracy was similar when decoding from DS only versus DS and Non-DS populations over short integration times (Fig 4F). There is a small decrease in error when including Non-DS cells when decoding with longer integration times (Fig 4F). Indeed, the OQE and OLE using DS cells alone provided similar accuracy (Fig 4G), indicating little benefit from this form of non-linear decoding in these conditions. Thus, decoding of the direction of motion continuously with short integration bins is effectively performed by a linear decoder that integrates signals from a large tuned population with little benefit from correlated activity between tuned or untuned neurons. Note, we are not saying the other classes of decoders, more explicitly constructed to decode motion from untuned neurons (e.g. Reichardt detectors to calculate direction of motion de novo) would not be useful for bolstering signals from DSGCs.

### Noise and temporal integration dictate spatial decoding constraints in model

The analyses above indicate that large populations of nearly independent ooDSGCs can be leveraged to rapidly decode the direction of motion, while temporal integration increases the importance of spatial correlation structure for accurate decoding. Ostensibly, the relationship between temporal integration and sensitivity to local correlations could be explained by the presence of high temporal frequency noise in ooDSGC responses. To better understand how noise and temporal integration influence population decoding we created a model that simulated responses from complete ooDSGC mosaics of various size and organization (see Methods). In brief, each modeled DS-unit response was generated from a distinct linear-nonlinear model (Fig 5A) with its position and direction orientation determined by one of four modeled mosaics (Fig 5B). The linear filter provides direction tuning and the non-linearity was adjusted to generate on-off responses with sparsity similar to that in the data. The DS-units were stimulated with a moving image used on the retina and the direction of motion was decoded at each time point from the population responses (Fig 5C). Note, decoding performance for the model (Fig 5C) was better than for the data (Fig 2C), because 532 simulated DS-units (133 for each direction) were used in the model, compared to just ~50 real DSGCs to decode with data. To test the spatial sensitivity of the decoder, populations were decoded either from local subsets (Fig 5Bii) or from DS-units with randomly chosen spatial locations (Fig 5Biii).

To begin, the response of the DS-units was noiseless. In this case, correlations between nearby DS-units was much higher than that observed in the measured data (Fig 5D, black curve). These correlations are caused by local image statistics and while they are diminished by the nonlinearity that produces the sparse responses, they remain very high between cells with high receptive field overlap. In the absence of noise, decoding error was substantially increased when decoding from DS-units with random spatial locations (Fig 5Ei) or shuffled responses (data not shown) – supporting previous work illustrating the importance of maintaining correlations in decoding direction (Franke et al., 2016; Zylberberg et al., 2016). Adding independent noise to each DS-unit reduced local correlations substantially (Fig 5D red curve) and greatly reduced the sensitivity of decoder performance on the spatial arrangement of the DS-units (Fig 5Eii). Finally, temporally integrating the noisy responses partially rescued the correlation structure established by the natural image (Fig 5D blue curve) and increased the sensitivity of the decoder to spatial structure (Fig 5Eiii). This model illustrates that temporal integration influences correlation structure (at least in this example) and that it also dictates the decoder’s reliance on those correlations. This helps to resolve the discrepancy between this study and previous studies which have highlighted the importance of correlations for decoding DSGC population responses: Here, brief temporal integration was required to decode dynamic global motion, while previous work focused long temporal integration because motion stimuli were local and had a static velocity.

## Discussion

To fully understand neural function, activity must be measured within an appropriate ecological context. While completely natural stimuli are difficult to produce and parameterize (Rust and Movshon, 2005), increasing stimulus complexity towards more natural contexts can help evaluate ideas about neural function derived from using simpler stimuli. In this paper, we assayed the potential of populations of ooDSGCs to signal the direction of dynamic global motion of a natural scene. While the stimulus we used falls short of capturing the full complexity of global motion as a mouse moves through the environment, it offers some features beyond drifting gratings or moving spots, while also providing more control over the stimulus than natural movies. For example, this stimulus has naturalistic statistics such as spatial structure that falls off (on average) as the inverse of the spatial frequency, but with high-contrast edges and ‘object’s not present in pink noise. Also, the frequency of direction changes can be manipulated to analyze how decoding performance depends on frequent direction changes (Fig 2D) versus constant velocity motion (Fig 2G).

Using this stimulus, we found that mammalian ooDSGCs integrate motion signals over ~200 ms and respond sparsely, but reliably over repeated presentations. This sparsity necessitates long integration times to decode accurately using small ooDSGC populations, which precludes decoding frequent changes in motion direction accurately and rapidly. On the other hand, large populations of ooDSGCs can be used to accurately and rapidly decode dynamic changes in the direction of global motion. Below, we discuss these findings in the context of previous research.

### Functional role of ooDSGCs

Several lines of evidence have supported the role of ooDSGCs in coding local motion. This is based primarily on three observations: 1) ooDSGCs exhibit diminished spike rates to global relative to local motion (so-called surround suppression) (Vaney et al., 2001; Chiao and Masland, 2003; Olveczky et al., 2003; Hoggarth et al., 2015); 2) oDSGCs do not exhibit surround suppression; 3) ooDSGCs have small receptive fields and a high density, which seem unnecessary for signaling global motion (Vaney et al., 2001).

Regarding ‘1’, recent work shows direction selectivity is maintained in global motion and that surround suppression may preferentially attenuate luminance responses (Hoggarth et al., 2015; Im and Fried, 2016). This suggests that the attenuation may emphasize direction information rather than obscuring it. Regarding ‘2’, oDSGCs exhibit similar response structure to ooDSGCs to dynamic global motion (Fig 1). While the incompleteness of our oDSGC populations prevented an analysis of decoding their responses, the similarity in their encoding properties to ooDSGCs suggests similar decoding performance. Regarding ‘3’, we showed that large dense populations are actually necessary to accurately and rapidly signal dynamic global motion given the sparsity of ooDSGC responses. Finally, a recent study showed that the cardinal axes formed by ooDSGCs align with the axes of self-motion on the retina, suggesting that the population is geared to signal global motion (Sabbah et al., 2017). Thus, we think it is plausible that ooDSGCs play a significant role in signaling global motion.

None of these arguments rule out a role for ooDSGCs to also encode local motion (see below). ooDSGCs clearly respond vigorously to moving spots and bars, which may be reasonable proxies for local motion in nature.

### Challenges and constraints to continuously decoding ooDSGC population responses

Response sparsity and stimulus variability preceding a spike (i.e. broad tuning) challenges a downstream decoder that must (nearly) continuously and accurately estimate a dynamic stimulus. This is a distinct decoding regime from that often investigated: i.e. when stimuli produce high firing rates (Theunissen and Miller, 1991; Fiscella et al., 2015; Marre et al., 2015; Kuhn and Gollisch, 2019) in a population of narrowly tuned neurons (Jazayeri and Movshon, 2006). Previous studies decoding dynamic motion stimuli from retina have utilized the optimal linear filter approach (Warland et al., 1997)}(Marre et al., 2015; Kuhn and Gollisch, 2019). This approach temporally integrates spike rates with a fixed filter to provide a continuous optimal linear estimate. Our approach differs from the optimal linear filter approach because we explored a range of integration times. Integrating over or under the optimal temporal range will increase the total mean squared error of the estimate but can decrease the error at specific temporal frequencies. Thus, this approach can highlight tradeoffs inherent in decoding visual information. While previous studies have used long temporal integration windows to improve decoding (Fiscella et al., 2015) we show that such strategies come at a significant cost to temporal resolution for decoding rapidly changing stimuli (Fig 2D).

To better illustrate the spatial-temporal tradeoff in decoding we constructed a simple model of signal and noise separation (Fig 6). In this model, noise is separated from signal using a simple linear filter that integrates over space and time (Fig 6A). As we observed for decoding ooDSGC population responses, temporal integration can improve signal detection by preferentially attenuating high frequency noise. This noise reduction strategy is limited though by the presence of a high (temporal) frequency signal, as further integration begins to degrade both signal and noise (Fig 6B_i_). The same problem occurs when integrating spatially (Fig 6B_ii_).

**Figure 6.**
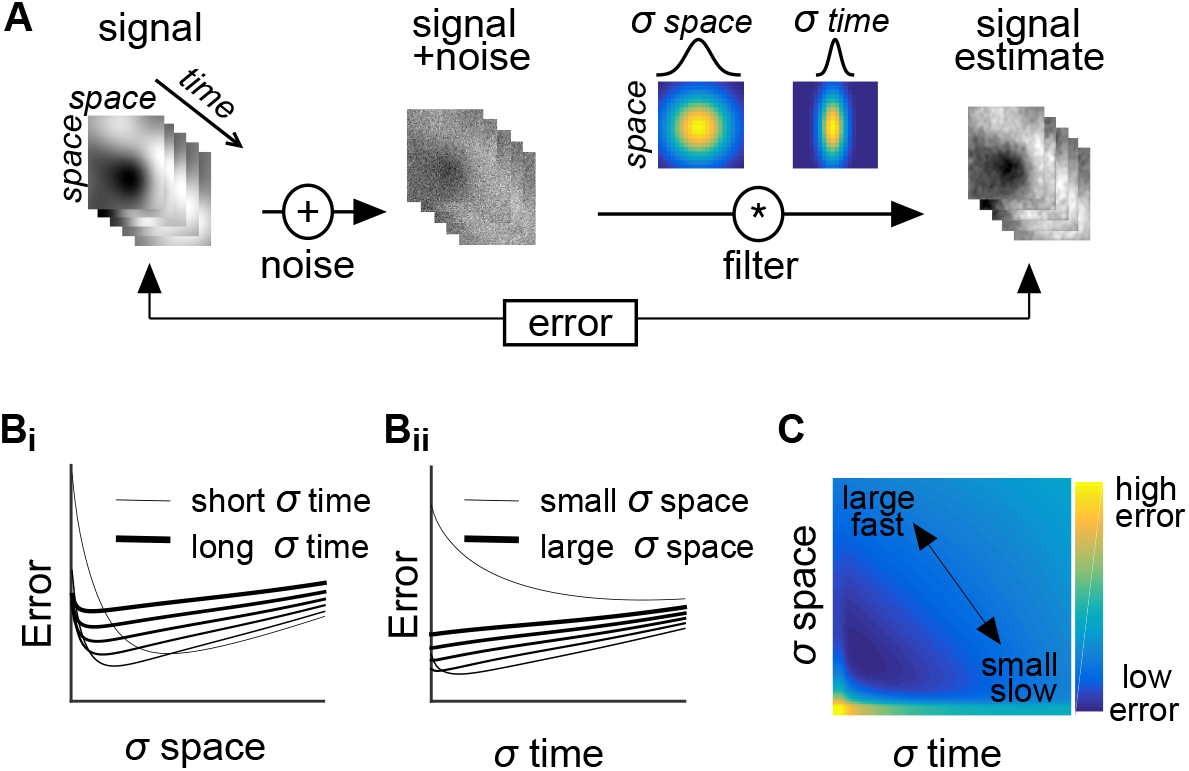
Separating signal and noise using spatial or temporal integration, sparing resolution in the dimension not integrated. **A.** Schematic of simple model illustrating signal corrupted by noise, integration with spatial and temporal linear filters, and estimate comparison. **B.** Error as a function of integrating spatially (Bi) or temporally (Bii) for signals that have been integrated with increasingly long temporal (Bi) or large spatial (Bii) filters. **C.** Error as a function of the temporal and spatial filter size.

However, the greater the spatial integration, the less temporal integration is needed to improve signal detection, sparing the high frequency temporal signal. A similar trade-off can be made in the opposite direction, by sacrificing temporal resolution for greater spatial resolution. Thus, in extracting information, a decoder can choose to focus on spatial or temporal resolution, at a cost to the temporal or spatial resolution, respectively (Fig 6C). Dynamic global motion requires high temporal resolution but minimal spatial resolution: We show that the population response of ooDSGCs permits a regime for accurately decoding this stimulus at short timescales with a simple linear decoder.

### Comparison to salamander DSGCs

This study complements a conceptually similar study recently performed in the salamander retina (Kuhn and Gollisch, 2019). In that study, textures were presented and dynamically displaced while recording responses from populations of OFF DSGCs. These OFF DSGCs may be analogous to mammalian oDSGCs because they exhibit minimal surround suppression and are organized along three directions (Kuhn and Gollisch, 2016). Like our study, the authors examined both encoding and decoding performance.

For *encoding*, salamander OFF DSGCs responded in a manner sensitive to the direction of global motion and integrated motion over ~200 ms, similar to the results presented here, It is interesting to note that the integration was similar despite the difference in species and the substantial differences in the recording temperatures: 21 °C (salamander) and 34°C (mouse).

For *decoding*, there are several differences between the two studies. First, we used the OLE (and OQE) while the previous study used the ‘optimal linear filter’ approach (see previous section for comment on these approaches) (Warland et al., 1997). Second, the previous study quantified decoding performance in terms of Shannon information, while we quantified it in terms of degrees of visual angle. Third, the previous study focused on the synergy available for decoding when cells with different preferred directions were used by the decoder, while we focused on how decoding performance depended on population size, decoding timescale, and correlation structure. Nevertheless, a few coherent comparisons can be made. First, both studies show that rapid changes in the direction of global motion are poorly decoded from small populations of DSGCs. Second, both studies indicate that noise correlations play a minimal role in rapidly decoding global motion.

Our study departs from previous work by analyzing global motion processing in the mouse retina. Furthermore, we explicitly show that (1) the encoding and response statistics of oDSGCs and ooDSGCs are similar for global motion; (2) large populations of ooDSGCs can be used to decode global motion with high temporal resolution; (3) the nature of ooDSGC population codes depends on the nature of the stimulus and the constraints of the decoding task, such as the necessity of high temporal versus high spatial resolution.

### The importance of correlation on neural decoding

The role of correlated activity in neural coding is intensely debated in neuroscience (Schneidman et al., 2003; Latham and Nirenberg, 2005; Averbeck et al., 2006; Shamir, 2014). Mammalian ooDSGCs have provided a useful model system to understand the sources and impact of correlated activity (Amthor et al., 2005; Franke et al., 2016; Zylberberg et al., 2016). Those studies largely pointed to correlations exerting a benefit upon DSGC population codes. We show that this result depends on context. When integrating (or decoding) over long timescales, correlation strength can be high and can improve decoding performance (Fig 4D). However, when integrating (or decoding) over short timescales, correlations are small, even over many cells, and thus decoding performance is independent of the correlations (Fig 4C-D). We also show that correlations between DSGC and non-DS RGC weakly impacted decoding error over short integration times (Fig 4G). These observations suggest that shared noise exists at lower temporal frequencies than independent noise. Thus, as demonstrated in a model (Fig 5), temporal integration diminishes independent noise and strengthens correlations in local populations, shifting the decoder’s input from independent to locally correlated populations.

These observations have implications for downstream circuits that process retinal signals. Circuits that decode local motion stimuli can leverage temporal integration to diminish independent noise without sacrificing spatial resolution. This favors a decoder utilizing local correlations, which help maintain a robust population response during a transient stimulus. In contrast, downstream circuits that decode global motion stimuli can pool over large numbers of ooDSGC assuming independence to achieve a nearly continuous readout of motion direction. This suggests distinct decoding regimes - decoders for large, fast visual stimuli relying on independent inputs and decoders for small and slow stimuli using correlated inputs. This may help explain why ooDSGC axons diverge to multiple downstream brain circuits including the lateral geniculate nucleus and superior colliculus (Kay et al., 2011). Future work may reveal that these distinct circuits instantiate these distinct decoding regimes. It is also possible that neuromodulators alter the integration times within a single circuit (Higley et al., 2009), switching between the two decoding regimes dynamically depending on current task demands.

## Acknowledgements

We would like to thank G. Awatramani, N-Y Jun, B. Murphy-Baum, L. Osborne, S. Roy, X. Yao, for comments on drafts of this manuscript. We thank T. Mrsic-Flogel, and F. Iacaruso for providing a movie from a camera attached to the head of a mouse. This work was supported by the Sloan Research Fellowship in Neuroscience (J.Z.), Canadian Institute for Advanced Research (CIFAR) (J.Z.), Azrieli Global Scholar Award for Learning in Machines and Brains (J.Z.); the Canada Research Chairs program (J.Z), the Whitehead Scholars Program at Duke University (G.D.F) and NEI/NIH grant R01 EY024567 (G.D.F.).

## Methods

### Mice and retina dissection procedures

Retinas were removed and recorded from C57BL/6J and CBA/CaJ x C57BL/6J mice between the ages of 1 month and 1 year. The strains showed no differences to the results reported in this study, thus data were pooled. Mice were used in accordance with the Duke University Institutional Animal Care and Use Committee.

Retina dissection was optimized to maintain response sensitivity. Mice were dark-adapted overnight, euthanized via decapitation, eyes were enucleated, and a piece of retina (~1-2mm^2^) was isolated from the pigmented epithelium (Yao et al., 2018). Retina isolation was performed in Ames solution (room temperature) bubbled with 95% O_2_ and 5% CO_2_. All procedures were performed in the dark under IR light. The retina was isolated from the dorsal half of the eye (identified from vasculature) to increase the fraction of M-opsin expressed in the cones for better overlap with the spectrum of the visual display.

### Multi-electrode array recording, spike sorting, cell position determination

Electrical activity was measured from RGCs on a multielectrode array (MEA). Spikes were identified, sorted into individual cell clusters, and soma positions on the MEA were estimated as previously described (Litke et al., 2004; Yao et al., 2018). Electrical activity was measured from RGCs using a hexagonal large-scale MEA, which was ~490 μm along an edge with 30 μm spacing between 519 electrodes (Field et al., 2010). Retinas were held against the MEA with a permeable membrane and were perfused with Ames solution (34 °C) bubbled with 95% O_2_ and 5% CO_2_.

Electrical activity was analyzed offline to identify and sort spikes into individual cell clusters (Shlens et al., 2006). Briefly, on each electrode, spikes were identified by a voltage threshold and voltage waveforms were concatenated across the six surrounding electrodes. These concatenated waveforms were parameterized with principal components analysis (PCA) and clustered with a mixture of gaussians model, providing putative cell assignments. Putative cells were analyzed if their spike time autocorrelation showed less than 10% refractory period violations and 25% spike time correlation with a cell identified on a nearby electrode, indicating spikes were from a single and uniquely identified neuron.

Soma position on the array was used to identify quadruplets (Fig 2) and pairwise distances between cells (Fig 3A). Soma position was estimated from the Electrical Image (EI) on the array (Li et al., 2015; Yao et al., 2018). The EI consisted of the average voltage on each electrode preceding a spike(Petrusca et al., 2005; Field et al., 2009). The position of the soma was taken as the center of mass of the EI.

### Visual stimulus

The retina was stimulated at photopic light levels (8000 Rh*/s) with a gamma-corrected OLED display (SVGA+XL Rev3 from eMagin). Three types of visual stimuli were presented to the retina and controlled via custom software written in Matlab utilizing the MGL library (gru.stanford.edu). First, drifting gratings, at two different temporal frequencies, were used to identify ooDSGCs (see RGC classification). Second, natural images, taken from the van Hateren image database (van Hateren and van der Schaaf, 1998), were presented to probe RGCs responses to global motion in natural images. Natural images were presented in two different stimulus protocols; using either dynamic or static velocity. Finally, natural movies from a head cam mounted mouse (from lab of Thomas Mrsic-Flogel) and cat (Betsch et al., 2004) were used to further test response sparsity.

In the dynamic velocity protocol, the same image was presented in the same orientation on every frame at image locations, X and Y. The average frame rate was 40 Hz (~25 ms/frame). X and Y were drawn randomly from a gaussian distribution and smoothed in time with a sliding window ~7.5 seconds. This generated an image that jittered around on the screen with a slow drift, presenting jitter across different image locations. X and Y were rectified to prevent displaying the image edge. The change in position between frames, ΔX(t) and ΔY(t), was sampled from an independent gaussian distributions with no temporal correlations (white). The standard deviation of the displacement distributions was ~20 um - corresponding to ~800 um/s along a single axis. A single dynamically moving image was presented for 60 minutes.

In the static direction protocol, the image was drifted in a single direction at ~1080 um/s for ~4 seconds before a new direction or image was presented. The image was reoriented for each direction with its longer edge parallel to the direction of motion. Six different images were presented at 8 different directions, spread equally across 360°.

### RGC classification

oDSGCs and ooDSGCs were identified based on their responses to square wave gratings (960 μm/cycle) drifted in 8 different directions and two different speeds (1 and ¼ Hz). Responses to each grating were quantified by total spike number generated during the 8 seconds each grating was presented. Cells were first clustered as DSGCs, then separated as ON-OFF and ON cells, and then grouped by their preferred direction (Yao et al., 2018). Direction-selective cells were clustered by their direction-selective indices (DSI) (Ravi et al., 2018) at each grating speed using a 2 dimensional gaussian mixture model. This method avoided setting an arbitrary threshold on DSI. Cells were then clustered by hand using the ratio of their response vector magnitudes for fast and slow gratings. Cells that maintained or increased their vector magnitudes for faster gratings were identified as ON-OFF. This process is based on the speed tuning curve differences between ON and ON-OFF DSGCs in mouse retina (Yao et al., 2018). Finally, ooDSGCs were clustered by hand based on the direction of their vector sum. Clustering by their preferred direction was only used to color code Figure 1C and did not contribute to decoding (see Optimal linear estimator).

### Optimal linear and optimal quadratic estimators

An optimal linear estimator (OLE) and optimal quadratic estimator (OQE) were used to estimate the direction of stimulus motion from a set of RGCs responses (Salinas and Abbott, 1994; Shamir and Sompolinsky, 2004). To do this, the OLE and OQE weight and sum the responses for each RGC:

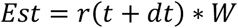

For the OLE RGC responses, r, were quantified by the number of spikes within a set number of sequential frames (bins) as indicated by the temporal integration time. *r* includes an added constant that allows for a default (offset) direction weight. For the OQE, *r* includes not only responses of individual cells but also the cross products of all possible cell pairs (Shamir and Sompolinsky, 2004). dt allows a time delay between response and the stimulus and was optimized to minimize the mean squared error between the direction estimate and true direction. The weights, W, were determined during a separate training set using Matlab’s backslash operation:

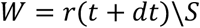

S is the cartesian coordinates of the stimulus direction. Matlab’s backslash operation returns a least-squares solution to a system of linear equations. Training sets for the dynamic motion stimulus consisted of the first ¾ time points and it was tested on the last ¼ time points. Training and testing on fully separated data blocks prevents the decoder from leveraging the PSTH autocorrelation to improve its test error. For the static motion stimulus, the OLE and OQE were trained on 5 images and tested on a hold-out image. Training and testing were redone for each image and errors averaged across all images.

To break correlation structures between cells, binned responses within the test data for the static direction stimulus were circularly shifted by a random amount independently for each cell. This manipulation maintained the direction selectivity of the response averages but broke correlations between cells.

### ooDSGC simulation

A simulation of ooDSGC receptive fields was used to test the observed results in larger and more complete mosaics than available from the measured data. The model consisted of four independent mosaics of modeled receptive fields responding to a moving image. As in the analysis of the measured data, the responses of the modeled neurons were combined to estimate the direction of image motion.

The response, R, of an individual model neuron was:

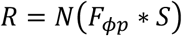

Where F_ϕp_, is the linear filter with preferred direction, ϕ, and position p. Linear filters consisted of a set of 10 sinewaves that were phase shifted 8 pix/frame, multiplied by a gaussian with a std of 17 pixels, centered at p and cut off in a square 75 pixel window. S is the stimulus; * indicates convolution; and N is a nonlinear function. The position, p, of each filter, F, was determined by an exclusion zone algorithm for generating 2 dimensional spatial mosaics (Galli-Resta et al., 1999). N was chosen as rectified linear function symmetric about zero – allowing responses to be ON-OFF with some control of the response sparsity. The threshold of the nonlinearity was adjusted to provide a similar fraction of spikes as that measured in the ooDSGCs population (90-95% of the bins had no activity). The density of the simulated mosaics was 30 cells/mm^2^ based on reported values in ooDSGCs in rabbit (Vaney, 1994). The population response, *<u>R</u>*, was decoded based on a linear model Est = *Σ*(*<u>R</u>*\*ϕ).

To understand how high frequency noise constrained decoding error we added gaussian white noise with a variance equal to the signal variance. Decoding was performed on the noisy signals without manipulation or after averaging from 10-frame bins.

### Spatial-temporal tradeoff simulation

A simulation of signal estimation was used to understand the tradeoffs inherent in separating signal from noise through spatial and temporal integration. The signal, S, was constructed by convolving spatial and temporal gaussian filters with white noise, creating a signal dominated by low spatial and temporal frequencies. Unfiltered white noise was then added to S. Then the noisy signal was filtered by convolving spatial and temporal Gaussian filters defined by their standard deviations, sigma. The mean squared error was calculated between the filtered output and S.

